# Characterizing Hydrogen Bonds in Intact RNA from MS2 Bacteriophage Using Solid State Magic Angle Spinning NMR

**DOI:** 10.1101/2021.06.02.446732

**Authors:** Orr Simon Lusky, Moran Meir, Amir Goldbourt

## Abstract

Ribonucleic acid (RNA) is a polymer with pivotal functions in many biological processes. RNA structure determination is thus a vital step towards understanding its function. The secondary structure of RNA is stabilized by hydrogen bonds formed between nucleotide base pairs and it defines the positions and shapes of functional stem-loops, internal loops, bulges, and other functional and structural elements. In this work we present a methodology for studying large intact RNA biomolecules using homonuclear ^15^N solid state nuclear magnetic resonance (NMR) spectroscopy. We show that Proton Driven Spin Diffusion (PDSD) experiments with long mixing times, up to 16s, improved by the incorporation of multiple rotor-synchronous ^1^H inversion pulses (termed Radiofrequency Dipolar Recoupling, RFDR, pulses), reveal key hydrogen-bond contacts. In the full-length RNA isolated from MS2 phage, we observed strong and dominant contributions of G-C Watson-Crick base pairs, and beyond these common interactions, we observe a significant contribution of the G-U wobble base pairs. Moreover, we can differentiate base-paired and non-base-paired nitrogen atoms. Using the improved technique facilitates characterization of hydrogen-bond types in intact large-scale RNA using solid-state NMR. It can be highly useful to guide secondary structure prediction techniques, and possibly structure determination methods.

## Introduction

RNA structure has long been a subject of intensive studies, mainly in view of the discovery that RNA is not only an information transfer molecule (mRNA) that transfers the DNA genetic code to the ribosome. Non-coding RNA molecules adopt a large variety of three-dimensional structures serving many cellular roles (1, 2). tRNA delivers amino acids to the ribosome that also contains ribosomal RNAs as part of its structure. Riboswitches regulate gene expression, ribozymes can catalyse various reactions similarly to enzymes, and additional roles of RNA exist and continue to be discovered, the most recent one being of RNA-glycan conjugates displayed on the cell surface (3).

Generally, most structural studies are performed on small synthetic RNA oligomers. For example, structures solved by NMR (4, 5) and X-ray crystallography (6) reveal various secondary structure elements such as base-paired helices, stem-loops, bulges and more. Structures of protein-RNA complexes are also prevalent, and were determined using various techniques such as cryogenic electron microscopy (CryoEM) (7), X-ray (8) and solution NMR (9). Solid state NMR has also provided structures of isolated RNA molecules or in complex with proteins (10, 11).

Large RNA molecules (beyond 100-200 nucleotides) are challenging for solution NMR due to high spectral overlap. For larger sizes, decreased relaxation times further complicate structure determination. Such limitations are partially solved by segmentally labelling the RNA to reduce the spectral congestion, or by using custom synthesis of nucleotides with partial enrichment (12, 13). RNA structures are also hard to obtain by crystallography techniques, and thus advanced sequence-based algorithms are a key tool to predict their secondary structure. Such techniques are based on minimizing the free energy on the basis of the hydrogen bond patterns (14, 15). Other methods for assessing the structure of large RNA molecules utilize enzymatic digestion, radical labelling and more (16).

The base pairs in polynucleic acids are stabilized by the hydrogen bonds. Canonical base pairs are formed between the nucleobases adenine (A) and uracil (U), and cytosine (C) and guanine (G). The canonical and most common Watson-Crick (WC) base-pairs are formed between the nucleobases, and in RNA are referred to as the ‘Watson-Crick edge’ (17). This type of bond is illustrated in figure 1A. However, there are other possible geometries for hydrogen bonds to form, namely the sugar edge and the Hoogsteen edge, as illustrated in figure 1B. Two nucleotides can form hydrogen bonds involving each pair of the three edges. Together with cis and trans orientations of the glycoside bond, there can be twelve types of hydrogen bonds between nucleotides giving the RNA flexibility in tertiary structure formation. Therefore, detection of the interacting edge of the hydrogen bond is essential to understanding RNA structure.

**Figure 1:**
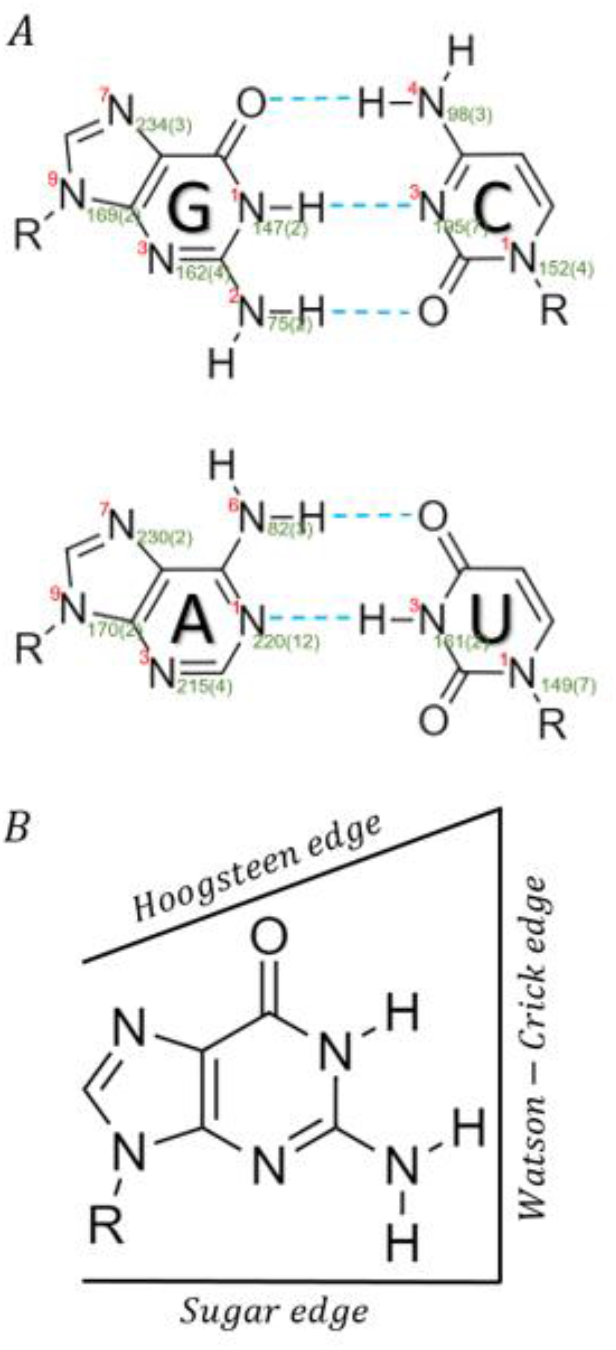
(A) Scheme of canonical Watson-Crick hydrogen bonds between nucleotides. The dashed blue lines represent the hydrogen bonds. BMRB nomenclature is in red and average ^15^N chemical shifts and standard deviations (in ppm) in green. R stands for ribose ring. (B) A guanine nitrogenous base with its three edges types.

Hydrogen bonds can be distinguished and characterized by NMR (18, 19) . In particular, ^2^J_NN_ scalar couplings between donor and acceptor nitrogen spins were observed using HNN-COSY solution NMR and report directly on hydrogen bonds in a 69 nucleotide-long RNA oligomer (20). For large RNA molecules that are highly challenging to solution studies, ^15^N spins across the hydrogen-bond are coupled by dipolar interactions that are not averaged in precipitated or sedimented samples and are thus amenable to solid-state NMR studies. Solid-state NMR has been successfully used to study various types of biological systems including folded and unfolded proteins, protein-DNA complexes, protein-RNA complexes, intact viruses and more (21–23). Recent progress in RNA characterization has been reviewed by Marchanka (24) and by Wang (25). Those mainly include synthetic RNA oligomers with or without bound proteins. For example, a 23-mer RNA fragment of HIV RNA and a (CUG)_97_ oligomer were studied using ^15^N-^15^N correlations (26, 27) and by ^1^H-detected ^15^N correlations at fast MAS (28). However, intact non-repetitive long RNA leads to high spectral congestion, which does not easily allow a resonance assignment for each nucleotide separately. Yet, the NMR correlations still hold information on the native hydrogen bonding patterns.

RNA viruses are common in nature. The size of their genome can be a few thousands of bases as in small spherical bacteriophages (29), or as large as 30 thousand bases, as in the case of SARS-CoV-2 (30). RNA viruses are fast mutating and therefore pose constant risk for global pandemics (31), as encountered in this past year and a half. The packing, folding, and capsid binding properties of RNA within the context of intact viruses is therefore important to understand and we constantly seek methods aimed to study intact viruses or isolated intact RNA.

MS2 bacteriophage infects *Escherichia Coli* (*E. Coli*) bacteria bearing positive F pili. It contains a 3,569 nucleotide-long single stranded RNA, encapsulated in a capsid made of 89 copies of coat protein dimers, and a maturation protein. Despite being the first RNA molecule to be sequenced (32), initially structural studies focused on interactions of the capsid protein with small RNA epitopes (33–35). Only recently the interaction of the full-length RNA with the MS2 capsid in a wild-type viral particle was studied in detail using CryoEM. Initially a resolution of 8.7Å (36) was obtained revealing a network of stem loop regions in the RNA. Later on, the resolution was significantly improved to 3.6Å for the capsid and 6Å for the RNA (37). This resolution was sufficient for tracing the secondary structure motifs of different segments in the RNA revealing unprecedented details on its structure. In particular, the identity of the bases involved in base-pair interactions could be determined for approximately two-thirds of the sequence.

Here we show how ^15^N correlations in solid-state NMR can be used to study the intact isolated RNA extracted from the MS2 phage. By improving ^15^N-^15^N proton-driven spin-diffusion polarization transfer (38) with rotor-synchronous π pulses applied to the ^1^H channel (as in radio-frequency driven recoupling (39)), which we term PDSD-RFDR, we show how hydrogen-bonds can be detected and characterized. From the type of nitrogen atoms involved, we can also estimate the type and face of those hydrogen bonds. We then compare the patterns we observe in the isolated RNA to that of the enclosed RNA derived from cryoEM.

## Materials and Methods

### Sample preparation

Wild type MS2 bacteriophage was produced by infecting cultures of *E. coli* strain C3000 and purified using our lab protocols for phage preparation (40). The RNA was harvested from the phage using a method similar to the one described by Meir et. al. (41). Phage were vortexed in TRIzol (TriReagent®), precipitated in isopropanol, followed by cold ethanol wash. The total yield of RNA from one litter bacterial culture was on average ~24 mg. The RNA precipitate was packed into a 4mm ZrO_2_ MAS rotor using a centrifuge and then used for NMR experiments. More details on MS2 phage preparation and purifications as well as RNA isolation appear in the supporting information (SI).

### NMR methods

Experiments were carried out on Bruker Avance III NMR spectrometers operating at 9.4T and 14.1T, both equipped with MAS 4mm probes. Two-dimensional ^15^N-^15^N correlation experiments were collected using PDSD (38) experiments with mixing times up to 16s. Additional PDSD experiments were acquired by adding rotor-synchronized π pulses (as in RFDR (39)) to the ^1^H channel during the mixing time (2s, 8s). We term this experiment PDSD-RFDR. A complete list of experimental parameters appears in the supporting information, Table S1. All experiments on the 9.4T magnet were performed at a spinning speed of 8 kHz and a set temperature of −28°C. The chemical shifts of ^15^N were externally referenced to ^15^NH_4_Cl at 39.3ppm (42).

### Data Analysis

NMR data were processed using TopSpin3.5 and NMRPipe (43). Analysis was performed using SPARKY version 3.134 (44).

### Numerical Simulation

The NMR simulation package SIMPSON (45) was used to verify the physical basis for the enhancement by RFDR but is not intended to reproduce experimental enhancements. A SIMPSON script was written to show that the application of π pulses on protons enhances the magnetization transfer between two X spins in a X_2_H_3_ spin system (X≡^13^C in this case, and represents a low-γ spin ½). The script, the dipolar couplings used, and the magnetization build-up curves appear in the SI.

## Results and Discussion

### Identification of ^15^N resonances

A typical two-dimensional ^15^N-^15^N proton-driven spin-diffusion (PDSD) solid-state MAS NMR correlation spectrum of MS2-RNA is shown in figure 2A. The off-diagonal correlation signals result from the dipolar coupling between ^15^N nuclei that are proximate in space, probably not beyond ~4 Å. In order to obtain them with sufficient sensitivity it was necessary to significantly increase the mixing time up to 16s (exploiting the long T_1_ of the ^15^N spins) and simultaneously reduce the spinning speed to 8 kHz (thus reducing MAS averaging of the dipolar interaction) while maintaining a set temperature of −28 °C. Similarly to our previous study on intact T7 bacteriophage dsDNA (46), we see clear asymmetry in the spectrum as well resulting from the excitation difference between primary, secondary, and tertiary amines, and between nucleobases having more or less protons.

**Figure 2:**
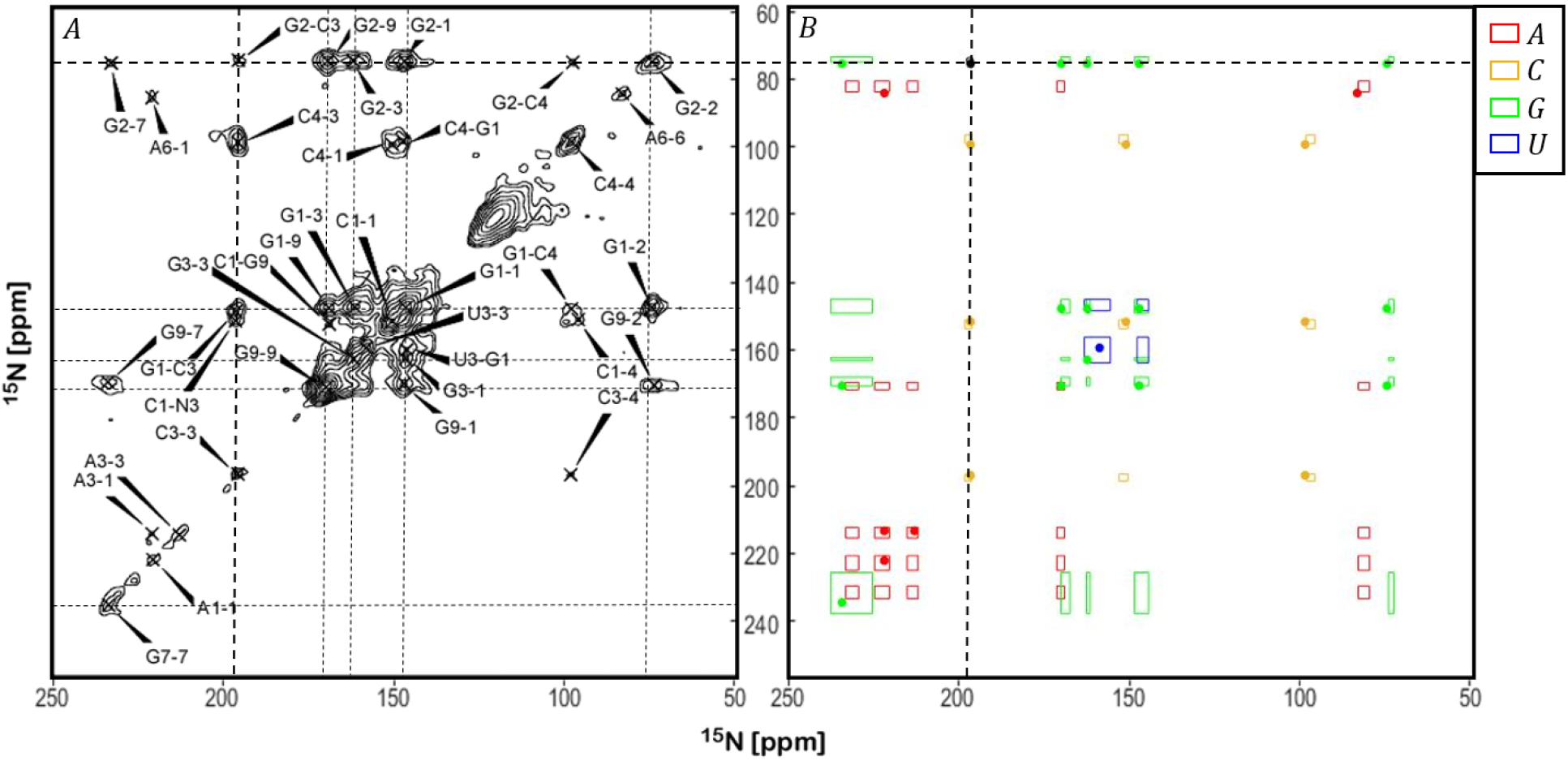
(A) A typical 2D ^15^N-^15^N PDSD spectrum of the MS2 RNA. A, C, G and U stand for the different nucleotides. Numbers identify the particular nitrogen in the nucleotide following BMRB nomenclature. Dashed lines mark the chemical shifts of G, and intersects between lines correspond to intra-nucleotide cross-peaks. The spectrum was acquired with a mixing time of 16s, a temperature set to −28°C, and spinning speed of 8 kHz. The spectrum was processed with an exponential line broadening of 100Hz in both dimensions. Twenty contours at multiples of 1.4 were generated with lowest contour set to a signal to noise (SNR) ratio of 5. (B) A theoretical plot of the most-probable positions of RNA intra-nucleotide signals. Coloured squares, different for each nucleotide, are centred around the average value of a particular peak according to a set of ten RNA oligomers, with their size corresponding to the standard deviations. Points mark the position of intra-nucleotide peaks observed in our spectra. The black ellipse represents an example of a hydrogen bond between G2 and C3 (see below). An equivalent plot that uses the average and standard deviation chemical shifts from the BMRB database is shown in figure S3 in the SI.

The resonances can be assigned to one of the four nucleobases shown in figure 1A but not to a specific nucleobase in the sequence (totalling 3569 bases). The assignment process relied on several strategies. Since BMRB averages for RNA oligomers have large standard deviations due to some errors in the database, initially data from ten representative RNA oligomers (see table S4 and figure S6 in the SI) was collected and their average chemical shifts and standard deviations were extracted (for comparison with BMRB see figure S5 in the SI). Figure 2B shows blocks centred at positions corresponding to correlations between those average RNA nucleobase shifts (differentiated by colour). Their size is given by the standard deviations of the shifts. Peaks from spectra similar to that in figure 2A were then marked on the predicted positions and assigned to the corresponding nucleotides. Many of the signals are easily identified. For example, primary amines are unique, having shifts of 82 ppm (Adenosine N6, or in short A6), 74 ppm (G2), and 97 ppm (C4). Additional unique individual shifts are C3 (197 ppm), A3 (214 ppm) and A1 (223 ppm). Other shifts have to be determined from two or three options (G1, U1 at ~146 ppm with C1 somewhat higher at 152; U3, G3 at ~160-162 ppm; G9, A9 at ~170 ppm; A7, G7 at ~231 ppm), with uracil being the most difficult to identify unambiguously. In order to resolve these ambiguities, we followed correlations to resolved signals performing “sidechain walks” (47) along proximate nitrogen nuclei in the same base. For example, in the spectrum shown in figure 2A the different ^15^N signals of the G-base are coloured and the linkage G2-G1-G9-G3-G7 can be clearly identified. Similarly, other nucleobases are assigned. Overall, we were able to identify all but one nitrogen shift, as shown in Table 1.

**Table 1:** ^15^N chemical shifts (in ppm) of MS2-RNA. The Unassigned nucleus A9is indicated by a grey cell. Dashes indicate non-existing atoms.

|  | N1 | N2 | N3 | N4 | N6 | N7 | N9 |
| --- | --- | --- | --- | --- | --- | --- | --- |
| A | 222.0 | - | 213.2 | - | 83.8 | 231.2 |  |
| C | 151.4 | - | 196.8 | 99.0 | - | - | - |
| G | 147.5 | 75.1 | 162.7 | - | - | 234.4 | 170.3 |
| U | 150.5 | - | 159.1 | - | - | - | - |

### Identification of hydrogen bonds

Once ^15^N shifts have been assigned, several inter-nucleotide contacts could be identified. Those are attributed almost exclusively to base-pairing interactions, which are characteristic of the helical double-strand-like secondary structure of the RNA (in MS2, 68% of the RNA that appears in the electron density maps is attributed to a base-pairing arrangement). In figure 2B (the schematics of the RNA spectrum) the position of a representative base-pairing cross-peak is given by the ellipse, correlating G2 and C3 belonging to a G-C canonical WC base-pair. Other contacts have been similarly identified.

A clear evidence for base-pairing interactions is shown in the different ^15^N-^15^N spectra shown in figure 3. We can identify G-C base-pairing via the contacts G1-C3 and G2-C3. We can also identify C4-G3, and C4-G1. Although the PDSD experiment is not sufficiently quantitative, the fact the G1-C3 is the strongest cross-peak suggests that these contacts are typical WC pairs. According to a CryoEM structure by Dai et. al. (37), analysis of RNA secondary structure elements shows that ~54% of the hydrogen bonds in the structure are formed between G and C. It coincides with the fact that most hydrogen bond correlations found in our spectra were between these two nucleotides. Additionally, a WC G-C base pair involves 5 proton spins available to mediated magnetization transfer, while other possible base pairs have less. Moreover, all spectra show higher intensities for guanine ^15^N signals, followed by cytosine and the weakest signals are of adenine. G and C also appear more frequently in the MS2 RNA. Another canonical base pair is expected between A and U, and mainly A1/6 and U3 assuming that the WC face is the most common arrangement. However, we could not detect such contacts. Both diagonal signals of A6 and A1 are relatively weak. This is not surprising given the fact that for adenine the closest proton polarization source to N1 is that on the carbon AC2, and only two additional protons are available (on N6 amine) to polarize the entire nucleobase. We observe even weaker A1-A6 and A1-A3 cross-peaks at long mixing times. Thus, the expected cross-peaks of A1-U3 at 222.0-159.1 ppm, or A6-U3 at 83.8-159.1 are probably too weak to detect at the current experimental conditions (uracil contributes only a single proton to the hydrogen bond). Given the fact that A-U base pairs are stabilized by two hydrogen-bonds (and 3 protons), while those of G-C by three hydrogen-bonds (and 5 protons), it is likely that the A-U pair is also more mobile reducing further the ^15^N-^15^N dipolar interaction. The weak signal on A6 in comparison to G2 and C4 (all primary amines) is another indication for enhanced dynamics in the adenine nucleobase.

**Figure 3:**
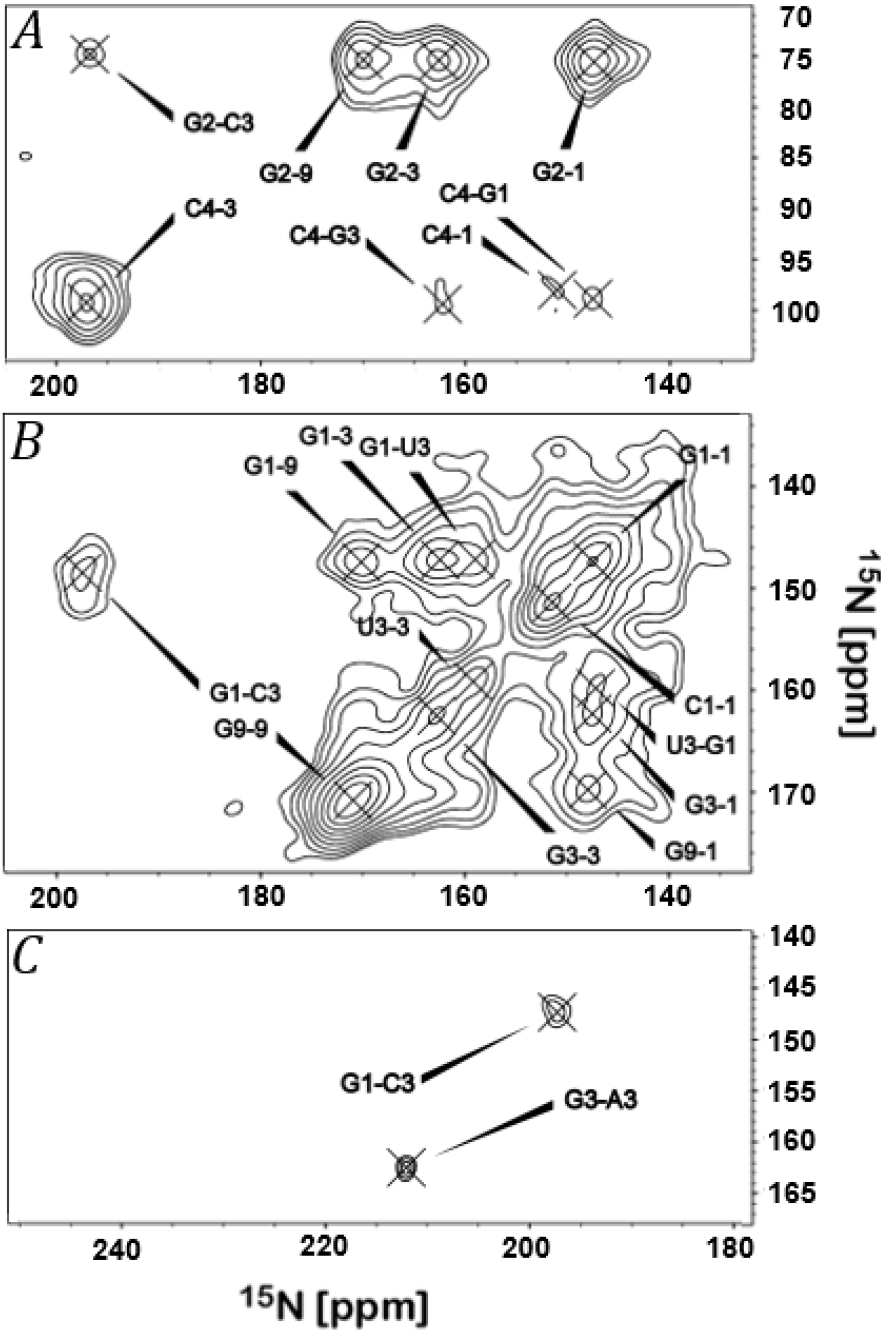
Examples of hydrogen-bond and additional inter-nucleotide cross-peaks from different ^15^N-^15^N spectra. The contour level was set such that the lowest signal has an SNR of 5. (A) PDSD-RFDR with a mixing time of 8s. (B) PDSD with a mixing time of 16s. This plot shows the G-U wobble basepair. (C) PDSD-RFDR, 2s.

In addition to the G-C canonical base pair, RNA structure is also based on the formation of G-U wobble base pairs (48). Interestingly, we observe (figure 3C) clear G1-U3 correlations between these two nucleotides, corresponding to the WC edge. Despite being energetically similar to the A-U base pair, the additional proton contributed by G, and its much preferred excitation efficiency provides sufficient polarization for this hydrogen-bond to be detected. The assignment to U3 of the peak at 159.1 ppm is based on the absence of any other correlations to G, where the G3 resonance at 162.3 is strongly correlated to G2 and G9. It is also supported by the correlation to U1 at 150.1 ppm, which we could detect using a PDSD spectrum taken at a higher field (figure S4 in the SI). According to the CryoEM structure, approximately 9% of the hydrogen bonds are formed between G and U.

Three additional uncommon contacts were observed in our spectra. Correlation peaks between G and A (G3-A1, G3-A3, G3-A6) could only be found in the PDSD-RFDR experiments. Hydrogen bonds between G and A were reported for bacterial ribosomal RNA, where A7 (part of the Hoogsteen edge) forms a bond with G1 (part of the WC edge) (49). However, in our case we observe correlations to G3, which is part of the sugar edge of the nucleobase. Another possibility is that we are detecting weak stacking interactions that correspond to the positioning of the N3 atoms at distances smaller then 4Å in space, or even closer if they belong to a curved region. Indeed, in MS2 PDB structures a few such contacts could be detected. A weak (SNR of 5) A6-C4 contact is also detected (figure 3A) in the PDSD-RFDR 8s spectrum. The low intensity can indicate a low abundance hydrogen bond, or more likely, another stacking interaction.

Another significant observation would have been the identification of signals belonging to non-base-paired nucleobases. Moreover, if the ratio of base-paired to non-base-paired could be determined quantitatively, the information could be input to structure prediction/calculation programs. According to the small set of RNA oligomers we summarized, shifts up to approximately 3 ppm or more are expected for some ^15^N spins. Others are not significantly affected. Observation of such shifts requires that simultaneously a pair of nitrogen spins will be involved in a hydrogen-bond and will be sufficiently shifted not to be obscured by the broadening of the signals resulting from averaging thousands of nucleotides. Figure 4 shows such an example for the cytosine nucleobase. Both C3 and C4 show broadening and shifting at the C3 diagonal signal around 197 ppm and at the C3-C4 cross-peak at 99 ppm. At the C3 position where the signals are most intense (197 ppm), also a base-pairing cross-peak G1-C3 can be detected (148 ppm). However, at the shoulders of the peaks, which have a higher chemical shift for C3 and a lower value for C4, no base-pairing cross-peak is detected. The reduced intensity of the shoulders and the absence of a cross-peak to guanine suggests that this signal belongs to non-base-paired cytosine nucleobases, existing at smaller percentages in the RNA structure.

**Figure 4:**
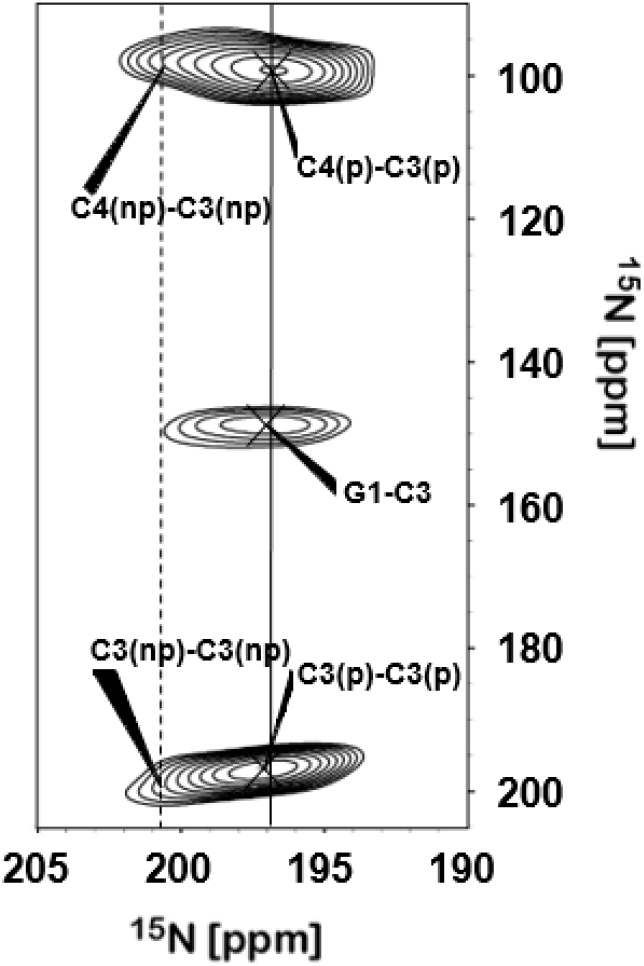
A segment from a 2D ^15^N-^15^N PDSD spectrum showing the distinction between base-paired (p) and non-base-paired (np) cytosine N3 and N4. The dash line indicates the shift of a non-base-paired C-N3. Solid line corresponds to base-paired C-N3.

At this stage the distinction of base-paired and non-base-paired nitrogen spins is qualitative, since RNA dynamics and the PDSD transfer mechanism do not allow a more quantitative information to predict the ratio of non-base-paired and base-paired contributions. For example, our spectra required temperatures below −25°C, above which hardly any signals could be detected at all. It is therefore possible that more dynamic RNA parts are still missing in our spectrum (with probably greater contribution of non-structured RNA regions) and that cross-peak signal intensities are affected by dynamics in addition to polarization transfer varying efficiencies. Future studies could address such points as was suggested for ^13^C spin-diffusion correlation experiments in proteins (50).

### Improving ^15^N polarization transfer with rotor-synchronous ^1^H π pulses

The homonuclear interaction between ^13^C spins is significantly stronger than that of ^15^N spin pairs and therefore short mixing times (10-500 ms) are sufficient for magnetization transfer. Moreover, by constantly irradiating on the ^1^H spins at a field that resonates with the spinning frequency (γB1=VR), known as dipolar assisted rotational resonance (DARR) (51), enhanced magnetization transfer is obtained. Yet, irradiating for seconds, as required for ^15^N recoupling, is in practice impossible due to hardware limitations, and therefore using DARR for ^15^N correlation experiments is not always feasible. Moreover, RNA is a dynamics biomolecule with a smaller density of proton spins as compared to proteins, further reducing the efficiency of dipolar recoupling. ^15^N correlation spectra in biological samples are therefore mostly obtained by the PDSD technique with the application of long mixing times (52, 53) in the order of seconds, or using direct polarization transfer via the application of rotor-synchronous ^15^N π pulses (26). At high spinning speeds, the PAR (proton assisted recoupling) technique has been proved useful (28, 54). PDSD is based on an indirect effect, in which the ^15^N-^15^N magnetization transfer is achieved due to ^1^H-^15^N dipolar recoupling (resulting from incomplete MAS averaging of this interaction).

Long mixing times required for ^15^N correlations always carry some signal decay due to relaxation and as the mixing times in our studies was increased from 2 to 16 seconds, the SNR was reduced by up to 7-fold in some cases (see Table S2 in the SI). On the other hand, new signals appeared. Although the homonuclear ^1^H-^1^H dipolar interaction is theoretically not necessary for the PDSD effect to occur (55), clearly enhanced ^1^H-^1^H interaction can increase the efficiency of ^1^H-^15^N heteronuclear dipolar recoupling. We therefore applied synchronous π pulses to the ^1^H channel following the RFDR scheme in order to directly recouple the ^1^H-^1^H homonuclear dipolar interaction. As demonstrated in figure S1 in the SI, this approach is different from the original suggestion to recouple directly heteronuclear interaction by applying two pulses every rotor period (55), or directly recoupling the homonuclear ^15^N dipolar interaction using synchronous π pulses (56).

In order to demonstrate this new approach, four ^15^N-^15^N correlation experiments were conducted – two with a mixing time of 2s, and two with a mixing time of 8s, each pair differing only by the application of the ^1^H-RFDR sequence. The overlay between the spectra is shown in figure 5. Although the average SNR at long mixing times is generally smaller, for most cross-peaks, the SNR has consistently increased (average of ~165% for the mixing time of 2s, and of ~127% for 8s) by the application of the π pulses. Moreover, not only does the PDSD-RFDR spectrum contain more correlations, these correlations are mostly between two hydrogen-bonded nitrogen spins thus providing essential additional information regarding the hydrogen bonds patterns in the RNA, enabling detection of the data described in previous sections. The actual SNR enhancement in both experiments are given in Tables S3a and S3b of the SI.

**Figure 5:**
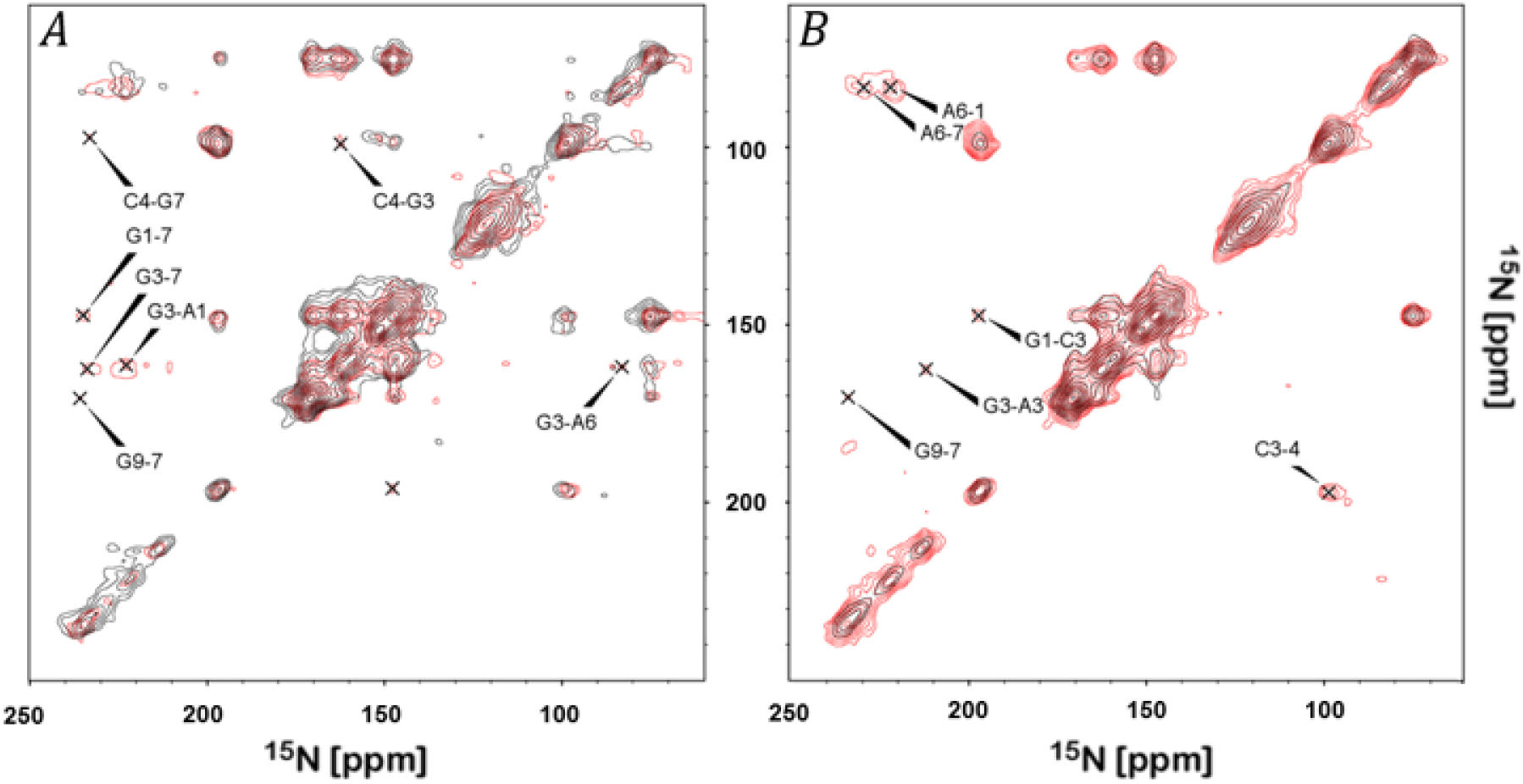
A comparison between PDSD (black) and PDSD-RFDR (red) ^15^N-^15^N correlation spectra acquired with a similar mixing time of (A) 8s and (B) 2s. Cross-peaks that appear only in the PDSD-RFDR experiment are assigned. Enhancement is also apparent in many other signals, averaging ~126% (8s) and ~153% (2s) for the cross-peaks. The contour levels were chosen such that the lowest SNR was 5. The processing (100Hz broadening, similar zero-filling, similar acquisition times) was identical in all cases.

In order to verify that the application of π pulses is the source of signal enhancement, we performed numerical simulations using the SIMPSON software (45). We calculated polarization transfer between two non-hydrogen (X) spins in a X_2_H_3_ spin system, where all spins are coupled by hetero- and homonuclear interactions. We followed in two cases the evolution of a preliminary state in which the two X spins are oppositely polarized; one with the π pulses (PDSD-RFDR), and another without. The numerical simulations shown in figure S2 in the SI show that indeed polarization transfer is enhanced with the π pulses. These results verify qualitatively our experimental results.

## Summary and Conclusions

The 1.1MDa full-length RNA isolated directly from the MS2 bacteriophage virus was studied using solid state NMR. Using RFDR-enhanced PDSD ^15^N-^15^N correlation experiments, we could assign the ^15^N resonances to the four different nucleotides, and detect hydrogen bonds, both canonical as well as wobble base pairs. These hydrogen bonds stabilize the secondary structure of the viral RNA and reduce its overall dynamics. By recognizing the ^15^N shifts making up the hydrogen bonds, it is possible with such techniques to determine the face of the bond and the dominant pairs we detect have a Watson-Crick face. Another observation is the ability to distinguish nitrogen spins involved in base-pairing interactions from those in non-base-paired nucleobases.

While we show how solid-state NMR can be useful to study such high-molecular-weight RNA biomolecules, detection of genomic ^15^N resonances in general, and more importantly hydrogen bonds, poses some challenges, some of which we have addressed here. Excitation of ^15^N signals is based on polarization transfer from ^1^H, and is therefore less efficient for tertiary amines and in particular in nucleotides bearing less proton spins. Thus guanine and cytosine signals dominate the spectral features. Another challenge is the ability to transfer polarization between ^15^N nuclei, which relies on the interaction with protons. While the standard techniques (at average spinning speeds) require extension of mixing times, the cost is signal decay due to T_1_ relaxation. Moreover, this increases significantly the total experimental time. We show here that by recoupling the ^1^H-^1^H dipolar interaction using radio-frequency driven-recoupling, signal enhancement is obtained without a need for seconds-long continuous irradiation that challenges the hardware. Consequently, the experimental time can be shortened significantly. It also allows gaining information even when long mixing times are inapplicable due to shorter relaxation times. Moreover, using this technique we have been able to detect new correlations not detected even at very long mixing times. Some of these correlations validate our assignment. Others are attributed to inter-nucleotide contacts. Towards more quantitative estimation of hydrogen bond patterns, it will be required to generate a more uniform and efficient ^15^N excitation, and estimate and fit polarization transfer between base pairs (57). The latter is required even if such hydrogen bonds are detected directly via proton detection (28).

The improved method we described for characterizing hydrogen bonds in intact RNA using solid-state NMR is applicable to RNA molecules extracted from natural sources, or even still embedded in their source organism, and can be utilized regardless of sequence length. Although not yet quantitative, it has the potential to guide structure prediction protocols.

## Acknowledgments

This research was supported by the Israel Science Foundation grant #847/17. We thank Prof. Uri Gophna from the Faculty of Life Sciences, Tel Aviv University, for supplying MS2 stocks, and for fruitful discussions.

## References

1. Eddy, S.R. (2001) Non-coding RNA genes and the modern RNA world. Nat. Rev. Genet., 2, 919–929.

2. Ransohoff, J.D., Wei, Y. and Khavari, P.A. (2018) The functions and unique features of long intergenic non-coding RNA. Nat. Rev. Mol. Cell Biol., 19, 143–157.

3. Flynn, R.A., Pedram, K., Malaker, S.A., Batista, P.J., Smith, B.A.H., Johnson, A.G., George, B.M., Majzoub, K., Villalta, P.W., Carette, J.E., et al. (2021) Small RNAs are modified with N-glycans and displayed on the surface of living cells. Cell, 10.1016/j.cell.2021.04.023.

4. Park, C.-J. (2003) Solution structure of the influenza A virus cRNA promoter: implications for differential recognition of viral promoter structures by RNA-dependent RNA polymerase. Nucleic Acids Res., 31, 2824–2832.

5. Barnwal, R.P., Yang, F. and Varani, G. (2017) Applications of NMR to structure determination of RNAs large and small. Arch. Biochem. Biophys., 628, 42–56.

6. Westhof, E. (2015) Twenty years of RNA crystallography. RNA, 21, 486–487.

7. Sugita, Y., Matsunami, H., Kawaoka, Y., Noda, T. and Wolf, M. (2018) Cryo-EM structure of the Ebola virus nucleoprotein-RNA complex at 3.6 Å resolution. Nature, 563, 137–140.

8. Ennifar, E., Nikulin, A., Tishchenko, S., Serganov, A., Nevskaya, N., Garber, M., Ehresmann, B., Ehresmann, C., Nikonov, S. and Dumas, P. (2000) The crystal structure of UUCG tetraloop 1 1Edited by J Doudna. J. Mol. Biol., 304, 35–42.

9. Wang, Z., Hartman, E., Roy, K., Chanfreau, G. and Feigon, J. (2011) Structure of a Yeast RNase III dsRBD Complex with a Noncanonical RNA Substrate Provides New Insights into Binding Specificity of dsRBDs. Structure, 19, 999–1010.

10. Ahmed, M., Marchanka, A. and Carlomagno, T. (2020) Structure of a Protein-RNA Complex by Solid-State NMR Spectroscopy. Angew. Chemie, 132, 6933–6940.

11. Marchanka, A., Simon, B., Althoff-Ospelt, G. and Carlomagno, T. (2015) RNA structure determination by solid-state NMR spectroscopy. Nat. Commun., 6, 7024.

12. Duss, O., Maris, C., von Schroetter, C. and Allain, F.H.-T. (2010) A fast, efficient and sequence-independent method for flexible multiple segmental isotope labeling of RNA using ribozyme and RNase H cleavage. Nucleic Acids Res., 38, e188–e188.

13. Marchanka, A., Kreutz, C. and Carlomagno, T. (2018) Isotope labeling for studying RNA by solid-state NMR spectroscopy. J. Biomol. NMR, 71, 151–164.

14. Turner, D.H., Sugimoto, N. and Freier, S.M. (1988) RNA Structure Prediction. Annu. Rev. Biophys. Biophys. Chem., 17, 167–192.

15. Reuter, J.S. and Mathews, D.H. (2010) RNAstructure: software for RNA secondary structure prediction and analysis. BMC Bioinformatics, 11, 129.

16. Strobel, E.J., Yu, A.M. and Lucks, J.B. (2018) High-throughput determination of RNA structures. Nat. Rev. Genet., 19, 615–634.

17. Leontis, N.B. and Westhof, E. (2001) Geometric nomenclature and classification of RNA base pairs. RNA, 7, S1355838201002515.

18. Flinders, J. and Dieckmann, T. (2006) NMR spectroscopy of ribonucleic acids. Prog. Nucl. Magn. Reson. Spectrosc.48, 137–159.

19. Grzesiek, S., Cordier, F., Jaravine, V. and Barfield, M. (2004) Insights into biomolecular hydrogen bonds from hydrogen bond scalar couplings. Prog. Nucl. Magn. Reson. Spectrosc., 45, 275–300.

20. Dingley, A.J. and Grzesiek, S. (1998) Direct Observation of Hydrogen Bonds in Nucleic Acid Base Pairs by Internucleotide 2 J NN Couplings. J. Am. Chem. Soc., 120, 8293–8297.

21. Ladizhansky, V. (2017) Applications of solid-state NMR to membrane proteins. Biochim. Biophys. Acta - Proteins Proteomics, 1865, 1577–1586.

22. Lecoq, L., Fogeron, M.-L., Meier, B.H., Nassal, M. and Böckmann, A. (2020) Solid-State NMR for Studying the Structure and Dynamics of Viral Assemblies. Viruses, 12, 1069.

23. Habenstein, B. and Loquet, A. (2016) Solid-state NMR: An emerging technique in structural biology of self-assemblies. Biophys. Chem., 210, 14–26.

24. Sreemantula, A.K. and Marchanka, A. (2020) Solid-state NMR spectroscopy for characterization of RNA and RNP complexes. Biochem. Soc. Trans., 48, 1077–1087.

25. Yang, Y. and Wang, S. (2018) RNA Characterization by Solid-State NMR Spectroscopy. Chem. - A Eur. J., 24, 8698–8707.

26. Leppert, J. (2004) Identification of NH...N hydrogen bonds by magic angle spinning solid state NMR in a double-stranded RNA associated with myotonic dystrophy. Nucleic Acids Res., 32, 1177–1183.

27. Riedel, K., Leppert, J., Ohlenschla, O. and Go, M. (2005) Characterisation of hydrogen bonding networks in RNAs via magic angle spinning solid state NMR spectroscopy. J. Biomol. NMR, 31, 331–336.

28. Yang, Y., Xiang, S., Liu, X., Pei, X., Wu, P., Gong, Q., Li, N., Baldus, M. and Wang, S. (2017) Proton-detected solid-state NMR detects the inter-nucleotide correlations and architecture of dimeric RNA in microcrystals. Chem. Commun., 53, 12886–12889.

29. Callanan, J., Stockdale, S.R., Shkoporov, A., Draper, L.A., Ross, R.P. and Hill, C. (2020) Expansion of known ssRNA phage genomes: From tens to over a thousand. Sci. Adv., 6, eaay5981.

30. Wu, F., Zhao, S., Yu, B., Chen, Y.-M., Wang, W., Song, Z.-G., Hu, Y., Tao, Z.-W., Tian, J.-H., Pei, Y.-Y., et al. (2020) A new coronavirus associated with human respiratory disease in China. Nature, 579, 265–269.

31. Carrasco-Hernandez, R., Jácome, R., López Vidal, Y. and Ponce de León, S. (2017) Are RNA Viruses Candidate Agents for the Next Global Pandemic? A Review. ILAR J., 58, 343–358.

32. Fiers, W., Contreras, R., Duerinck, F., Haegeman, G., Iserentant, D., Merregaert, J., Min Jou, W., Molemans, F., Raeymaekers, A., Van den Berghe, A., et al. (1976) Complete nucleotide sequence of bacteriophage MS2 RNA: primary and secondary structure of the replicase gene. Nature, 260, 500–507.

33. Sugiyama, T., Hebert, R.R. and Hartman, K.A. (1967) Ribonucleoprotein complexes formed between bacteriophage MS2 RNA and MS2 Protein in vitro. J. Mol. Biol., 25, 455–463.

34. Valegård, K., Murray, J.B., Stonehouse, N.J., van den Worm, S., Stockley, P.G. and Liljas, L. (1997) The three-dimensional structures of two complexes between recombinant MS2 capsids and RNA operator fragments reveal sequence-specific protein-RNA interactions. J. Mol. Biol., 270, 724–738.

35. Helgstrand, C. (2002) Investigating the structural basis of purine specificity in the structures of MS2 coat protein RNA translational operator hairpins. Nucleic Acids Res., 30, 2678–2685.

36. Koning, R.I., Gomez-Blanco, J., Akopjana, I., Vargas, J., Kazaks, A., Tars, K., Carazo, J.M. and Koster, A.J. (2016) Asymmetric cryo-EM reconstruction of phage MS2 reveals genome structure in situ. Nat. Commun., 7, 12524.

37. Dai, X., Li, Z., Lai, M., Shu, S., Du, Y., Zhou, Z.H. and Sun, R. (2017) In situ structures of the genome and genome-delivery apparatus in a single-stranded RNA virus. Nature, 541, 112–116.

38. Szeverenyi, N.M., Sullivan, M.J. and Maciel, G.E. (1982) Observation of spin exchange by two-dimensional fourier transform 13C cross polarization-magic-angle spinning. J. Magn. Reson., 47, 462–475.

39. Bennett, A.E., Griffin, R.G., Ok, J.H. and Vega, S. (1992) Chemical shift correlation spectroscopy in rotating solids: Radio frequency-driven dipolar recoupling and longitudinal exchange. J. Chem. Phys., 96, 8624.

40. Morag, O., Sgourakis, N.G., Abramov, G. and Goldbourt, A. (2018) Filamentous Bacteriophage Viruses: Preparation, Magic-Angle Spinning Solid-State NMR Experiments, and Structure Determination. In Methods in Molecular Biology. Vol. 1688, pp. 67–97.

41. Meir, M., Harel, N., Miller, D., Gelbart, M., Eldar, A., Gophna, U. and Stern, A. (2020) Competition between social cheater viruses is driven by mechanistically different cheating strategies. Sci. Adv., 6, eabb7990.

42. Bertani, P., Raya, J. and Bechinger, B. (2014) 15N chemical shift referencing in solid state NMR. Solid State Nucl. Magn. Reson., 61-62, 15–18.

43. Delaglio, F., Grzesiek, S., Vuister, G., Zhu, G., Pfeifer, J. and Bax, A. (1995) NMRPipe: A multidimensional spectral processing system based on UNIX pipes. J. Biomol. NMR, 6, 277–293.

44. Lee, W., Tonelli, M. and Markley, J.L. (2015) NMRFAM-SPARKY: enhanced software for biomolecular NMR spectroscopy. Bioinformatics, 31, 1325–1327.

45. Bak, M., Rasmussen, J.T. and Nielsen, N.C. (2000) SIMPSON: A General Simulation Program for Solid-State NMR Spectroscopy. J. Magn. Reson., 147, 296–330.

46. Abramov, G. and Goldbourt, A. (2014) Nucleotide-type chemical shift assignment of the encapsulated 40 kbp dsDNA in intact bacteriophage T7 by MAS solid-state NMR. J. Biomol. NMR, 59, 219–230.

47. Higman, V.A. (2018) Solid-state MAS NMR resonance assignment methods for proteins. Prog. Nucl. Magn. Reson. Spectrosc., 106-107, 37–65.

48. Varani, G. and McClain, W.H. (2000) The G·U wobble base pair. EMBO Rep., 1, 18–23.

49. Traub, W. and Sussman, J.L. (1982) Adenine-guanine base pairing in ribosomal RNA. Nucleic Acids Res., 10, 2701–2708.

50. Duong, N.T., Raran-Kurussi, S., Nishiyama, Y. and Agarwal, V. (2018) Quantitative 1 H-1 H Distances in Protonated Solids by Frequency-Selective Recoupling at Fast Magic Angle Spinning NMR. J. Phys. Chem. Lett., 9, 5948–5954.

51. Takegoshi, K., Nakamura, S. and Terao, T. (2001) 13C-1H dipolar-assisted rotational resonance in magic-angle spinning NMR. Chem. Phys. Lett., 344, 631–637.

52. Giraud, N., Blackledge, M., Böckmann, A. and Emsley, L. (2007) The influence of nitrogen-15 proton-driven spin diffusion on the measurement of nitrogen-15 longitudinal relaxation times. J. Magn. Reson., 184, 51–61.

53. Traaseth, N.J., Gopinath, T. and Veglia, G. (2010) On the Performance of Spin Diffusion NMR Techniques in Oriented Solids: Prospects for Resonance Assignments and Distance Measurements from Separated Local Field Experiments. J. Phys. Chem. B, 114, 13872–13880.

54. Lewandowski, J.R., Paëpe, G. De, Eddy, M.T. and Griffin, R.G. (2009) 15 N-15 N Proton Assisted Recoupling in Magic Angle Spinning NMR. J. Am. Chem. Soc., 131, 5769–5776.

55. Takegoshi, K., Nakamura, S. and Terao, T. (2003) 13C-1H dipolar-driven 13C-13C recoupling without 13C rf irradiation in nuclear magnetic resonance of rotating solids. J. Chem. Phys., 118, 2325–2341.

56. Robyr, P., Meier, B.H. and Ernst, R.R. (1989) Radio-frequency-driven nuclear spin diffusion in solids. Chem. Phys. Lett., 162, 417–423.

57. Duong, N.T., Raran-Kurussi, S., Nishiyama, Y. and Agarwal, V. (2020) Can proton-proton recoupling in fully protonated solids provide quantitative, selective and efficient polarization transfer? J. Magn. Reson., 317, 106777.

