## Supplementary material for "Characterizing Hydrogen Bonds in Intact RNA from MS2 Bacteriophage Using Solid State Magic Angle Spinning NMR": SI

**MS2 preparation**: A single colony of the host was grown in LB media until log phase (Optical density at 600nm of 0.6), and then introduced to a minimal media based on the M9 solution (1) (0.478M $Na_2HPO_4$, 0.220M $KH_2PO_4$, 0.086M NaCl ,2mM $MgSO_4$, 20μM $CaCl_2$, 4.04μM $FeSO_4$, 2μg Thiamine Hydrochloride). The medium also included 0.16g of $^{15}NH_4Cl$ as the sole nitrogen source, 1g of glycerol and 1g of sodium bicarbonate, as carbon sources.

The media was incubated at 37°C and shaken at 220rpm. Upon reaching $OD_{600}$ of 0.6, the shaking rate was reduced to 60rpm for an hour, allowing the host cells to grow the F pilus. Afterwards, the host was infected with MS2 stock in a multiplicity of ~100. The culture was incubated at 37°C and shaken at 220rpm until the $OD_{600}$ stabilized for at least two hours.

Bacteria cells were pelleted at 8000rpm at 4°C for an hour by centrifugation, the supernatant containing the phages was decanted and then the phages were precipitated by adding Polyethylene glycol (PEG) 8000 and NaCl to final concentrations of 100(W/V) and 1M respectively. The precipitate was pelleted at 10,000rpm at 4°C for 2 hours, and resuspended in 10mM TRIS pH=8 buffer. The solution was brought to a density of ~1.41 g/ml using CsCl (~6.8g CsCl for 10.75g solution) in an Ultra-Clear 13.2ml Beckman tubes and spun for 48 hours at a speed of 37000 rpm at 4°C in an Ultracentrifuge Optima XE 100K using an SW-41 Ti swinging bucket rotor. The band containing MS2 phage was collected from the tube, precipitated using PEG8000 in order to eliminate excess Cs ions, and resuspended in TRIS buffer PH=8.

**RNA extraction:**

MS2 phages in the TRIS buffer were precipitated using PEG 8000 and 1M NaCl, and pelleted by centrifugation at 10,000rpm at 4°C for two hours. The MS2 pellet was then resuspended in standard PBS solution, and aliquots of 1ml were transferred into RNase free Eppendorf tubes.

1ml of TriReagent was added to each tube, and the tubes were vortexed for 5 minutes. Afterwards, the tubes were centrifuged at room temperature for 5min at 13,000rpm. The upper band was collected into an RNase free Eppendorf, and an equal volume of isopropanol was added. The tubes were kept at -20°C overnight.

Subsequently, MS2-RNA was pelleted by centrifuge at 13,000rpm at 4°C for 20 minutes. The supernatant was discarded, and 1ml of cold ethanol (70%, RNase free) was added. After another centrifugation at the conditions above, the supernatant was discarded again, and the pellet containing full-length MS2-RNA was packed in a 4mm $ZrO_2$ MAS NMR rotor (~50μl).

**Figure S1** – PDSD-RFDR pulse sequence with π pulses applied synchronously to the $^1$H channel.

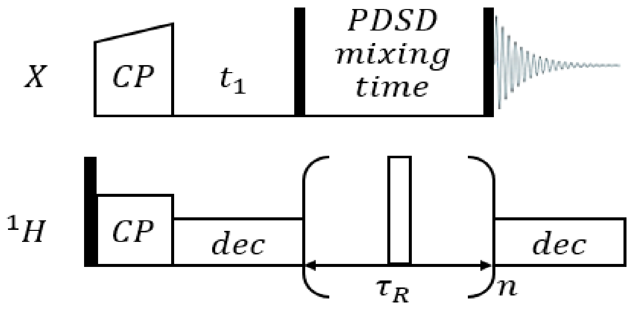


**Figure S2** – Simpson simulations demonstrating enhancement of polarization transfer by the addition of π pulses, an approach identical to the RFDR recoupling scheme. The system containing to low-γ spins ($^{13}$C in this case) and three protons. The simulations follow the magnetization of an inverted $^{13}$C spin $<I_{2z}>$ due to polarization transfer from a paired $^{13}$C spin in the presence of couplings to three homonuclear-coupled protons. The initial state of the density matrix was $I_{1z} - I_{2z}$ and is not normalized, $< I_{2z} > (0) = -8$. The script is given below.

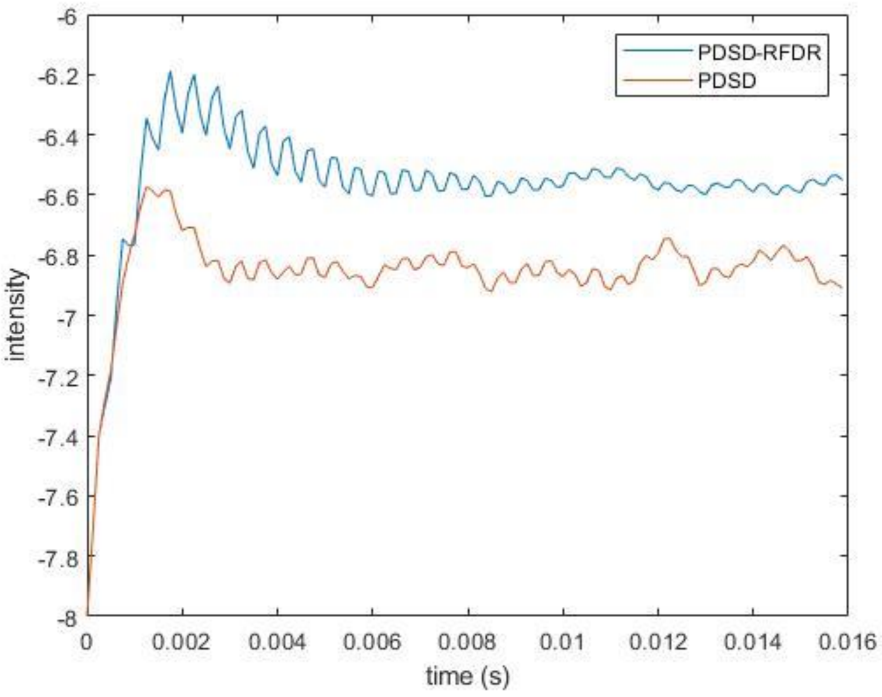

Simpson script for PDSD-RFDR:

```
spinsys {
    channels 13C 1H
    nuclei 13C 13C 1H 1H 1H
    shift 1 -1000 2p 0.0 -29.6 91.2 -19.4
    shift 2 5000 10p 0.0  29.6 21.2 -19.4
    dipole 1 2 -2000 0 0 0
    dipole 1 3 -21940 0 10 0
    dipole 1 4 -21940 0 70 0
    dipole 1 5 -21940 0 70 0
    dipole 2 3 -21940 0 10 0
    dipole 2 4 -21940 0 30 0
    dipole 2 5 -21940 0 30 0
    dipole 3 4 -25040 0 50 0
    dipole 3 5 -25040 0 150 0
    dipole 4 5 -55040 0 70 0
}
par {
    start_operator   I1z-I2z
    detect_operator  I2z
    spin_rate        8000
    gamma_angles     20
    sw               spin_rate
    crystal_file     rep100
    np               128
    verbose          1101
    proton_frequency 400e6
    conjugate_fid true
}
proc pulseq {} {
   global par
   set tr [expr 1.0e6/$par(spin_rate)]
   set tr2 [expr 0.5e6/$par(spin_rate)]

delay $tr2
delay $tr2
pulseid 5 0000 0 100000 0
store 1
reset
   for {set i 0} {$i<$par(np)    } {incr i} {
   acq
   prop 1
   }
}
proc main {} {
    global par
    set f [fsimpson]
    fsave $f $par(name).fid -xreim
}
```

**Table S1**: Experimental parameters

**Table S1a**: Parameters for homonuclear 2D $^{15}N$-$^{15}N$ PDSD experiments

| Experiment number | 1,2,3 | 4 | 5 |
|---|---|---|---|
| Field [T] | 9.4 | | 14.1 |
| Spinning frequency ($\nu_R$) [kHz] | 8 | | 13 |
| Set Temperature (°C) | -28 | | -30 |
| Acquisition points (t1/t2) | 196/496 | 220/1492 | 128/2994 |
| Acquisition time [ms] (t1/t2) | 9.80/9.92 | 11.0/29.84 | 2.56/29.94 |
| Carrier frequency [ppm] | 122.0 | | 122.8 |
| H 90° Pulse length [µs] | 3 | | 3.5 |
| N 90° Pulse length [µs] | 6 | | 4 |
| CP power ($\nu_H$) [kHz] | 83 | | 66 |
| CP power ($\nu_N$) [kHz] | 42 | | 55 |
| CP contact time [ms] | 1.5 | | 1.5 |
| PDSD mixing time [s] | 2,4,16 | 8 | 4 |
| $^{1}H$ decoupling power [kHz] | 83 | | 71 |
| swf-tppm decoupling pulse [µs] | 6 | | |
| Relaxation delay [s] | 4 | | |
| Scans | 64 | | 16 |
| SW F1/F2 [kHz] | 10/25 | | 50/25 |
| **Processing parameters F1/F2** | | | |
| Processing software | NMRPipe | | |
| Zero fill (t1/t2) | 16384/4096 | | |
| Apodization | Various exponential line broadenings (50-200 Hz) | | Exponential line broadening (100/250Hz) |

**Table S1b**: Parameters for homonuclear 2D $^{15}N$-$^{15}N$ PDSD-RFDR experiments

| Experiment number | 6 | 7 |
|---|---|---|
| Field [T] | 9.4 | |
| Spinning frequency ($\nu_R$) [kHz] | 8 | |
| Set Temperature (°C) | -28 | |
| Acquisition points (t1/t2) | 196/496 | 220/1492 |
| Acquisition time [ms] (t1/t2) | 9.80/9.92 | 11.0/29.84 |
| Carrier frequency [ppm] | 122.0 | |
| H 90° Pulse length [µs] | 3 | |
| N 90° Pulse length [µs] | 6 | |
| CP power ($\nu_H$) [kHz] | 83 | |
| CP power ($\nu_N$) [kHz] | 42 | |
| CP contact time [ms] | 1.5 | |
| PDSD mixing time [s] | 2 | 8 |

| | |
|---|---|
| $^1$H decoupling power [kHz] | 83 |
| swf-tppm decoupling pulse [µs] | 6 |
| Relaxation delay [s] | 4 |
| Scans | 64 |
| SW F1/F2 [kHz] | 10/25 |
| **Processing parameters F1/F2** | |
| Processing software | NMRPipe |
| Zero fill (t1/t2) | 16384/4096 |
| Apodization | Various exponential line broadenings (50-200 Hz) |

**Table S2:** Comparison of signal-to-noise ratio (SNR) between PDSD spectra acquired with different mixing times. All spectra were processed identically (100Hz broadening). Shaded area indicates lack of signal.

| Correlation | PDSD 2s | PDSD 4s | PDSD 8s | PDSD 16s |
|---|---|---|---|---|
| A1-1 | 15 | 12 | 11 | 8 |
| A3-1 | | | | 6 |
| A3-3 | 14 | 12 | 9 | 8 |
| A6-1 | | 6 | 7 | 7 |
| A6-6 | 24 | 15 | 11 | 8 |
| A6-G9 | | | | 6 |
| A7-7 | 17 | 14 | 14 | 8 |
| C1-1 | 90 | 84 | 92 | 86 |
| C1-4 | | | 6 | 8 |
| C3-3 | 22 | 16 | 15 | 10 |
| C3-4 | | 7 | 7 | 6 |
| C4-1 | | | 6 | 12 |
| C4-3 | 13 | 14 | 18 | 16 |
| C4-4 | 47 | 34 | 28 | 17 |
| C4-G1 | 8 | | 7 | 12 |
| G1-C3 | | | 7 | 12 |
| G1-1 | 114 | 92 | 93 | 81 |
| G1-2 | 17 | 14 | 20 | 17 |
| G1-3 | 24 | 22 | 33 | 34 |
| G1-9 | | 6 | 16 | 20 |
| G1-U3 | 18 | 16 | 25 | 26 |
| G2-C3 | | | 7 | 8 |
| G2-C4 | | | | 7 |
| G2-1 | 20 | 16 | 19 | 21 |
| G2-2 | 53 | 30 | 26 | 15 |
| G2-3 | 12 | 9 | 13 | 16 |
| G2-7 | | | | 7 |
| G2-9 | 8 | | 12 | 20 |
| G3-1 | 21 | 17 | 23 | 23 |
| G3-2 | | | 6 | |

| | | | | |
|---|---|---|---|---|
| G3-3 | 56 | 42 | 40 | 31 |
| G7-7 | 21 | 19 | 18 | 9 |
| G9-1 | 9 | | 5 | 17 |
| G9-2 | | | 6 | 11 |
| G9-7 | | | | 10 |
| G9-9 | 124 | 120 | 121 | 110 |
| U3-G1 | 20 | 15 | 20 | 22 |
| U3-3 | 47 | 36 | 33 | 25 |

**Table S3:** Comparison of SNR between PDSD and PDSD-RFDR acquired with similar mixing times (a: 2 sec, b: 8 sec). Cells coloured in grey correspond to cross-peaks that do not appear in a spectrum (SNR threshold of 5). Green cells correspond to correlations that have a better SNR in the PDSD-RFDR. Dagger marks cross-peaks that appear in the PDSD-RFDR spectrum but not in the PDSD spectrum. All spectra were processed with a line broadening of 100Hz.

**Table S3a:** Comparison of SNR for PDSD and PDSD-RFDR acquired with a mixing time of 2 sec.

| Correlation | PDSD | PDSD-RFDR | Difference |
|---|---|---|---|
| A1-1 | 15 | 24 | 160% |
| A3-3 | 14 | 27 | 193% |
| A6-1 | | 11 | † |
| A6-6 | 24 | 43 | 179% |
| A6-7 | | 8 | † |
| A7-7 | 17 | 31 | 182% |
| C1-1 | 90 | 153 | 170% |
| C3-3 | 22 | 26 | 118% |
| C3-4 | | 12 | † |
| C4-3 | 13 | 40 | 308% |
| C4-4 | 47 | 67 | 143% |
| G1-C3 | | 6 | † |
| G1-1 | 114 | 202 | 177% |
| G1-2 | 17 | 26 | 153% |
| G1-3 | 24 | 27 | 112% |
| G1-9 | | 8 | † |
| G1-U3 | 18 | 19 | 106% |
| G2-1 | 20 | 30 | 150% |
| G2-2 | 53 | 65 | 123% |
| G2-3 | 12 | 28 | 233% |
| G2-9 | 8 | 12 | 150% |
| G3-A3 | | 7 | † |
| G3-1 | 21 | 17 | 81% |
| G3-3 | 56 | 94 | 168% |
| G7-7 | 21 | 37 | 176% |
| G9-1 | 9 | | |
| G9-7 | | 6 | † |

| | | | |
|---|---|---|---|
| G9-9 | 124 | 202 | 163% |
| U3-G1 | 20 | 17 | 85% |
| U3-3 | 47 | 84 | 179% |

**Table S3b:** Comparison of SNR for PDSD and PDSD-RFDR acquired with a mixing time of 8 sec.

| Correlation | PDSD | PDSD-RFDR | Difference |
|---|---|---|---|
| A1-1 | 11 | 7 | 64% |
| A3-3 | 9 | 10 | 111% |
| A6-1 | 7 | 9 | 129% |
| A6-6 | 11 | 12 | 109% |
| A7-7 | 14 | 9 | 64% |
| C1-1 | 92 | 104 | 113% |
| C1-4 | 6 | 5 | 83% |
| C3-3 | 15 | 11 | 73% |
| C3-4 | 7 | 10 | 143% |
| C3-G1 | | 6 | † |
| C4-1 | 6 | 5 | 83% |
| C4-3 | 18 | 29 | 161% |
| C4-4 | 28 | 28 | 0% |
| C4-G1 | 7 | 6 | 86% |
| C4-G3 | | 6 | † |
| C4-G7 | | 5 | † |
| G1-C3 | 7 | 11 | 157% |
| G1-C4 | 8 | 8 | 0% |
| G1-1 | 93 | 107 | 115% |
| G1-2 | 20 | 21 | 105% |
| G1-3 | 33 | 35 | 106% |
| G1-7 | | 7 | † |
| G1-9 | 16 | 16 | 0 |
| G1-U3 | 25 | 17 | 68% |
| G2-C3 | 7 | 8 | 114% |
| G2-1 | 19 | 27 | 142% |
| G2-2 | 26 | 23 | 88% |
| G2-3 | 13 | 22 | 169% |
| G2-9 | 12 | 21 | 175% |
| G3-A1 | | 8 | † |
| G3-A6 | | 6 | † |
| G3-1 | 23 | 26 | 113% |
| G3-2 | 6 | 7 | 117% |
| G3-3 | 40 | 47 | 118% |
| G3-7 | | 6 | † |
| G7-7 | 18 | 14 | 78% |
| G9-1 | 5 | 11 | 220% |
| G9-2 | 6 | 8 | 133% |

| | | | |
|---|---|---|---|
| G9-7 | | 5 | † |
| G9-9 | 121 | 119 | 98% |
| U3-G1 | 20 | 18 | 90% |
| U3-3 | 33 | 35 | 106% |

**Figure S3 -** (A) A 2D $^{15}N$-$^{15}N$ PDSD spectrum of the MS2 RNA. This spectrum is identical to Fig. 2 in the article. A, C, G and U stand for the different nucleotides. Numbers identify the particular nitrogen in the nucleotide following BMRB nomenclature. Dashed lines mark the chemical shifts of G and intersects between lines correspond to intra-nucleotide cross-peaks. The spectrum was acquired with a mixing time of 16s at a spinning speed of 8 kHz, and a temperature set to -28°C. The spectrum was processed with an exponential line broadening of 100Hz in both dimensions. Twenty contours at multiples of 1.4 were generated with lowest contour set to a signal to noise ratio of 5. (B) A theoretical plot of the most-probable position of RNA intra-nucleotide signals according to the BMRB with their size corresponding to the standard deviation. Coloured squares, different for each nucleotide, are centred around the average value of a particular peak. Points mark the position of intra-nucleotide peaks observed in our spectra. The black ellipse represents an example of a hydrogen bond between G2 and C3.

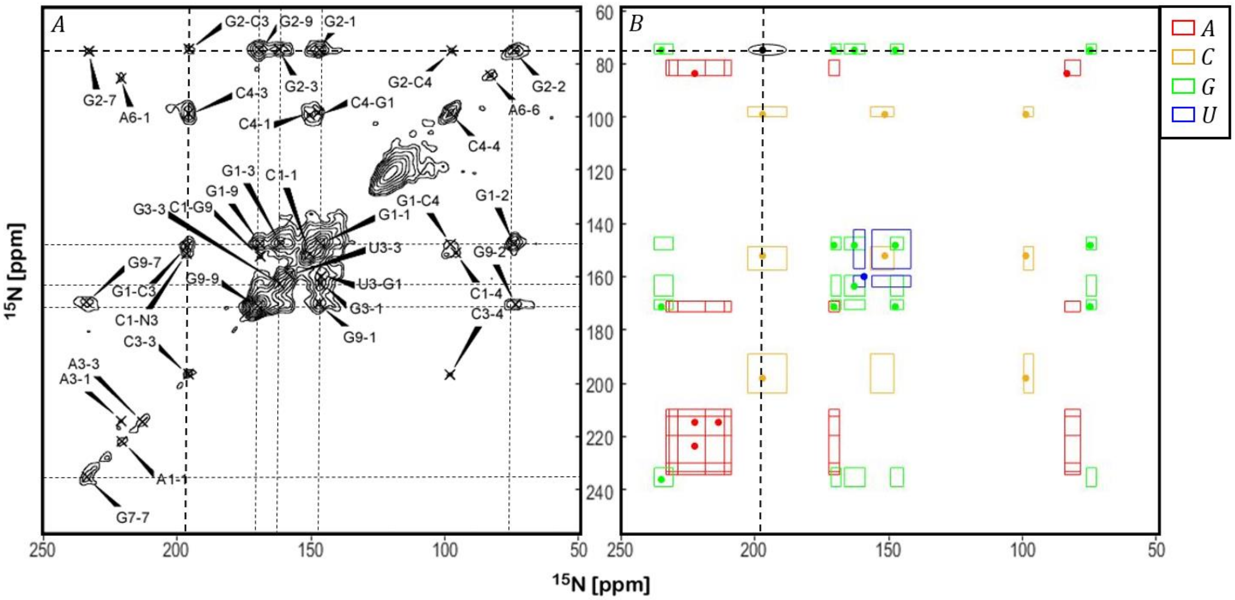

**Figure S4** - A 2D $^{15}N$-$^{15}N$ PDSD spectrum of the MS2 RNA acquired at a field of 14.1T. This spectrum resolves the small ambiguity between G and U and clearly identifies the signal corresponding to UN1 via the U1-3 cross-peak. Numbers identify the particular nitrogen in the nucleotides G and U following BMRB nomenclature. The spectrum was acquired with a mixing time of 4s at a spinning speed of 13 kHz, and a temperature set to -28°C. The spectrum was processed with an exponential line broadening of 100Hz and 200Hz in the direct and the indirect dimensions respectively. Twenty contours at multiples of 1.25 were generated with lowest contour set to a signal to noise ratio of 5.

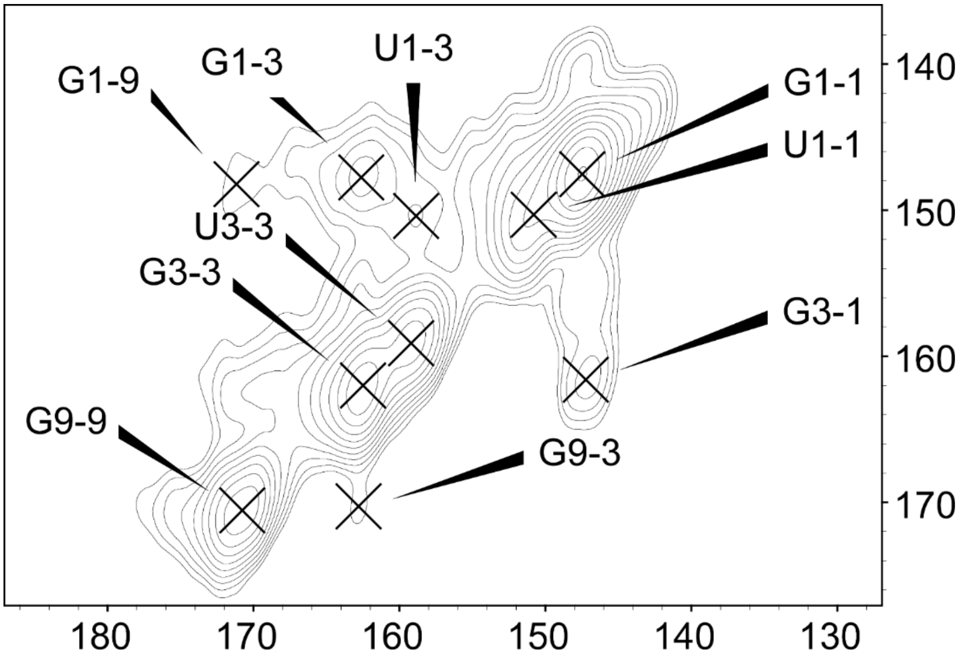

**Table S4: Average $^{15}N$ chemical shifts of 10 representative RNA oligomers.**

**Table S4a:** Average and standard deviation. The atom numbers follow the BMRB nomenclature.

| Nucleobase | $^{15}N$ atom number | Average chemical shift [ppm] | Standard deviation [ppm] | Counts |
|---|---|---|---|---|
| A | 1 | 222.71 | 2.17 | 23 |
| | 3 | 213.74 | 1.59 | 17 |
| | 6 | 81.82 | 1.63 | 7 |
| | 7 | 231.41 | 1.90 | 17 |
| | 9 | 170.44 | 1.11 | 22 |
| C | 1 | 152.11 | 1.35 | 29 |
| | 3 | 197.45 | 1.02 | 17 |
| | 4 | 97.40 | 1.29 | 18 |
| G | 1 | 146.83 | 2.05 | 58 |
| | 2 | 73.87 | 0.79 | 7 |
| | 3 | 162.40 | 0.46 | 6 |
| | 7 | 230.96 | 6.69 | 36 |
| | 9 | 169.06 | 1.33 | 47 |
| U | 1 | 146.44 | 1.66 | 22 |
| | 3 | 159.74 | 3.81 | 40 |

**Table S4b:** Sources of selected RNA oligomers

| **Source** | **BMRB entry** | **Number of nucleotides** | **Number of resonances** |
|---|---|---|---|
| Moloney Murine Leukaemia Virus U5-primer-binding-site (2) | 25049 | 68 | 26 |
| Rous Sarcoma Virus negative regulator of splicing (3) | 6062 | 23 | 14 |
| Human telomerase RNA CR7 terminal hairpin (4) | 7403 | 24 | 54 |
| | 7405 | 18 | 10 |
| Yeast C/D box small nucleolar RNA (5) | 34321 | 31 | 77 |
| HIV-1 Exon splicing Silencer 3 (6) | 17671 | 27 | 16 |
| HIV-1 transactivation-response element (7) | 11014 | 16 | 9 |
| Enterovirus internal ribosome entry site (8) | 6076 | 34 | 32 |
| HIV-1 frame-shift inducing site stem-loop (9) | 5834 | 22 | 31 |
| T2 bacteriophage gene 32 mRNA pseudoknot (10) | 4253 | 36 | 93 |

**Figure S5** - Comparison of the average chemical shifts, and their standard deviations, between the filtered BMRB data and the smaller set of 10 RNA oligomers indicated in Table S4a and shown in Figure S6.

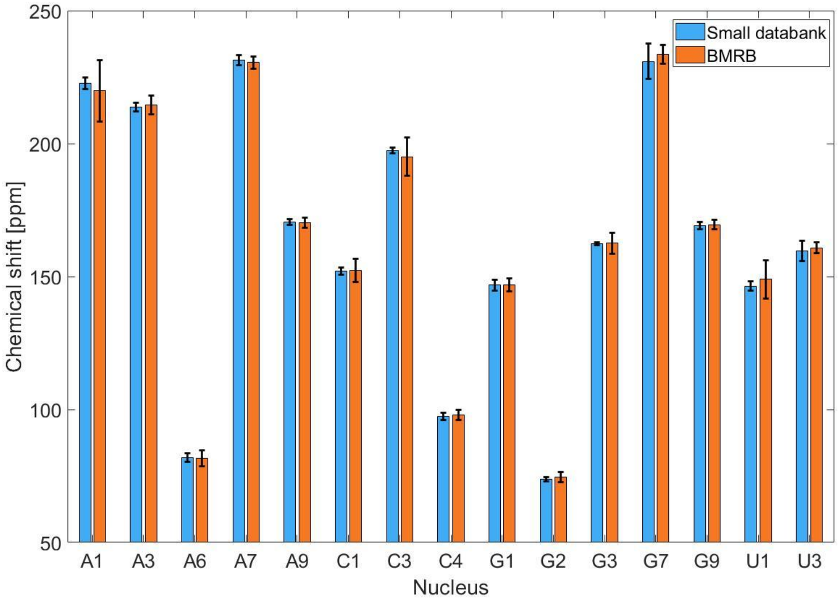


**Figure S6** - Secondary structure of the RNA oligomers that were used for generating an alternative database. The plots were made using RNApdbee (11, 12). Green lines represent first order pseudoknot interactions.

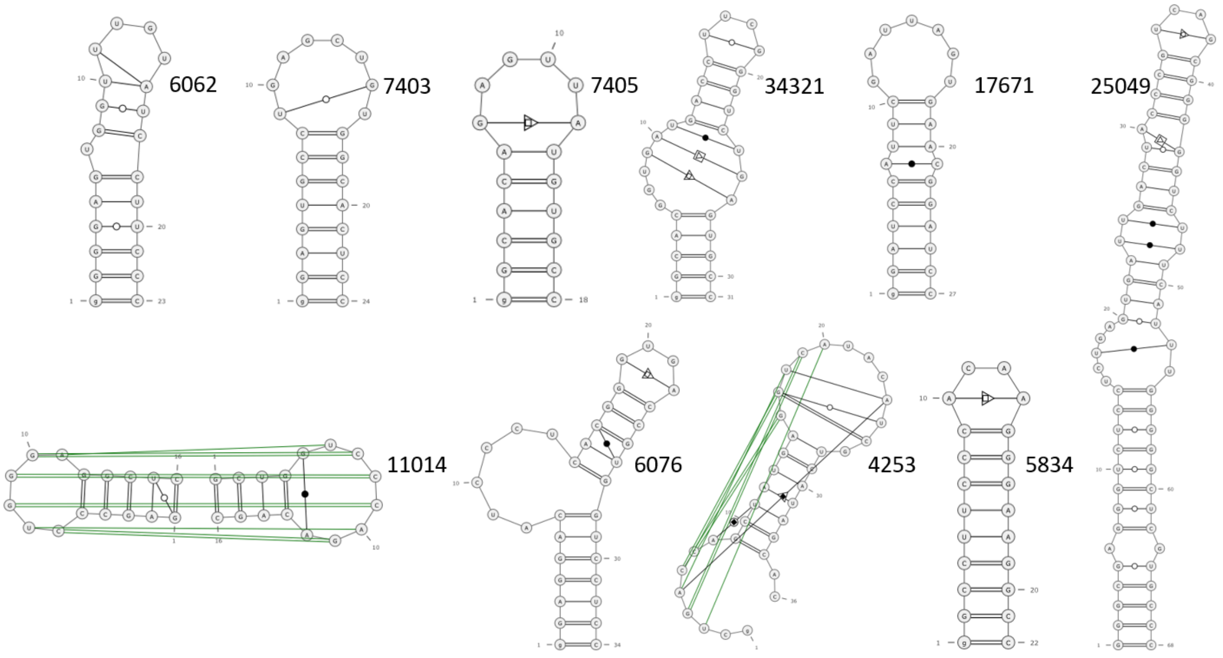

| RNA base-base classification | Visualization symbol |
|---|---|
| cis Watson-Crick Watson-Crick | ● |
| trans Watson-Crick Watson-Crick | ○ |
| cis Watson-Crick Hoogsteen | ■⃝ |
| trans Watson-Crick Hoogsteen | □⃝ |
| cis Watson-Crick Sugar | ▶⃝ |
| trans Watson-Crick Sugar | ▷⃝ |
| cis Hoogsteen Watson-Crick | ◙ |
| trans Hoogsteen Watson-Crick | ⧇ |
| cis Hoogsteen Hoogsteen | ■ |
| trans Hoogsteen Hoogsteen | □ |
| cis Hoogsteen Sugar | ▶⃞ |
| trans Hoogsteen Sugar | ▷⃞ |
| cis Sugar Watson-Crick | ●▶ |
| trans Sugar Watson-Crick | ○▷ |
| cis Sugar Hoogsteen | ■▶ |
| trans Sugar Hoogsteen | □▷ |
| cis Sugar Sugar | ▶ |
| trans Sugar Sugar | ▷ |

### Supporting information bibliography


1. Sambrook,J. and Russel,D.W. (2001) Molecular Cloning: A Laboratory Manual.

2. Miller,S.B., Yildiz,F.Z., Lo,J.A., Wang,B. and D'Souza,V.M. (2014) A structure-based mechanism for tRNA and retroviral RNA remodelling during primer annealing. *Nature*, **515**, 591–595.

3. Cabello-Villegas,J., Giles,K.E., Soto,A.M., Yu,P., Mougin,A., Beemon,K.L. and Wang,Y.-X. (2004) Solution structure of the pseudo-5' splice site of a retroviral splicing suppressor. *RNA*, **10**, 1388–1398.

4. Theimer,C.A., Jády,B.E., Chim,N., Richard,P., Breece,K.E., Kiss,T. and Feigon,J. (2007) Structural and Functional Characterization of Human Telomerase RNA Processing and Cajal Body Localization Signals. *Mol. Cell*, **27**, 869–881.

5. Chagot,M.-E., Quinternet,M., Rothé,B., Charpentier,B., Coutant,J., Manival,X. and Lebars,I. (2019) The yeast C/D box snoRNA U14 adopts a "weak" K-turn like conformation recognized by the Snu13 core protein in solution. *Biochimie*, **164**, 70–82.

6. Levengood,J.D., Rollins,C., Mishler,C.H.J., Johnson,C.A., Miner,G., Rajan,P., Znosko,B.M. and Tolbert,B.S. (2012) Solution Structure of the HIV-1 Exon Splicing Silencer 3. *J. Mol. Biol.*, **415**, 680–698.

7. Van Melckebeke,H., Devany,M., Di Primo,C., Beaurain,F., Toulme,J.-J., Bryce,D.L. and Boisbouvier,J. (2008) Liquid-crystal NMR structure of HIV TAR RNA bound to its SELEX RNA aptamer reveals the origins of the high stability of the complex. *Proc. Natl. Acad. Sci.*, **105**, 9210–9215.

8. Du,Z., Ulyanov,N.B., Yu,J., Andino,R. and James,T.L. (2004) NMR Structures of Loop B RNAs from the Stem−Loop IV Domain of the Enterovirus Internal Ribosome Entry Site: A Single C to U Substitution Drastically Changes the Shape and Flexibility of RNA ,. *Biochemistry*, **43**, 5757–5771.

9. Staple,D.W. (2003) Solution structure of the HIV-1 frameshift inducing stem-loop RNA. *Nucleic Acids Res.*, **31**, 4326–4331.

10. Holland,J.A., Hansen,M.R., Du,Z. and Hoffman,D.W. (1999) An examination of coaxial

stacking of helical stems in a pseudoknot motif: The gene 32 messenger RNA pseudoknot of bacteriophage T2. *RNA*, **5**, S1355838299981360.

11. Antczak,M., Zok,T., Popenda,M., Lukasiak,P., Adamiak,R.W., Blazewicz,J. and Szachniuk,M. (2014) RNApdbee—a webserver to derive secondary structures from pdb files of knotted and unknotted RNAs. *Nucleic Acids Res.*, **42**, W368–W372.

12. Zok,T., Antczak,M., Zurkowski,M., Popenda,M., Blazewicz,J., Adamiak,R.W. and Szachniuk,M. (2018) RNApdbee 2.0: multifunctional tool for RNA structure annotation. *Nucleic Acids Res.*, **46**, W30–W35.
